# Seascape genomics reveals metapopulation connectivity network of *Paramuricea biscaya* in the northern Gulf of Mexico

**DOI:** 10.1101/2021.10.06.463359

**Authors:** Matthew P. Galaska, Guangpeng Liu, Destiny West, Katie Erickson, Andrea Quattrini, Annalisa Bracco, Santiago Herrera

## Abstract

The degree of connectivity among populations influences their ability to respond to natural and anthropogenic stressors. In marine systems, determining the scale, rate, and directionality of larval dispersal is therefore central to understanding how coral metapopulations are interconnected and the degree of resiliency in the event of a localized disturbance. Understanding these source-sink dynamics is essential to guide restoration efforts and for the study of ecology and evolution in the ocean. The patterns and mechanisms of connectivity in the deep-sea (> 200 meters deep) are largely understudied. In this study, we investigated the spatial diversity patterns and metapopulation connectivity of the octocoral *Paramuricea biscaya* throughout the northern Gulf of Mexico (GoM). *Paramuricea biscaya* is one of the most abundant corals on the lower continental slope (between 1200 and 2500 m) in the GoM. The 2010 Deepwater Horizon oil spill (DWH) directly impacted populations of this species and thus are considered primary targets for restoration. We used a combination of seascape genomic analyses, high-resolution ocean circulation modeling, and larval dispersal simulations to quantify the degree of population structuring and connectivity among *P. biscaya* populations. Evidence supports the hypotheses that the genetic diversity of *P. biscaya* is predominantly structured by depth, and that larval dispersal among connected populations is asymmetric due to dominant ocean circulation patterns. Our results suggest that there are intermediate unsampled populations in the central GoM that serve as stepping stones for dispersal. The data suggest that the DeSoto Canyon area, and possibly the West Florida Escarpment, critically act as sources of larvae for areas impacted by the DWH oil spill in the Mississippi Canyon. This work illustrates that the management of deep-sea marine protected areas should incorporate knowledge of connectivity networks and depth-dependent processes throughout the water column.

## 1. Introduction

Marine ecosystems have traditionally been considered “open” with few apparent barriers to dispersal. However, phylogeographic studies often reveal unexpected levels of population structuring or even previously unrecognized cases of cryptic speciation (Hellberg, 2009; Hoffman et al., 2012; Cerca et al., 2021). These studies have primarily focused on coastal ecosystems and species of significant economic importance. In comparison, the patterns and mechanisms that generate genetic diversity in the deep-sea (> 200 m deep) are largely understudied (Baco et al., 2016; Taylor and Roterman, 2017).

One general pattern in the deep-sea is that populations found at different depths (vertically separated by tens to hundreds of meters) are generally more differentiated than populations found at similar depths over large geographical areas (horizontally separated by hundreds to thousands of kilometers) (Taylor and Roterman, 2017). However, the mechanisms responsible for this pattern remain poorly understood. Determining the scales of connectivity of marine populations and the mechanisms behind them is crucial for the conservation of marine ecosystems (Palumbi, 2003; Kinlan et al., 2005; Botsford et al., 2009; Gaines et al., 2010), and the study of diversification and evolution in the ocean (McClain and Mincks Hardy, 2010).

Population genetic methods enable the identification of genetic structuring patterns and estimate the scale, rate, and direction of reproductive exchange among marine populations (Breusing et al., 2016; Galaska et al., 2017; Bertola et al., 2020). These inferences, when coupled with analyses of environmental parameters, physical models of ocean circulation, and simulations of larval dispersal, can significantly enhance our understanding of connectivity networks at scales relevant to management (Benestan et al., 2016; Sandoval-Castillo et al., 2018; Xuereb et al., 2018; Bernatchez et al., 2019; Bracco et al., 2019). This integrative approach is known as seascape genetics (Galindo et al., 2006; Selkoe et al., 2016). Only a handful of studies have implemented seascape approaches in the deep sea. These have predicted the presence of intermediate “phantom” populations of hydrothermal vent species along mid-ocean ridges (Breusing et al., 2016) and have suggested that variables related to currents and food sources may explain a significant fraction of observed genetic patterns of sponge and coral species (Zeng et al., 2020).

Corals are essential foundational species in deep-sea benthic habitats and are typically slow-growing and long-lived (Roark et al., 2009; Sherwood and Edinger, 2009; Prouty et al., 2011, 2016; Girard et al., 2019). Deep-sea coral ecosystems are analogous to islands in that they are discrete and spatially separated. Each community serves as an oasis or biodiversity hotspot by locally enhancing the abundance and diversity of invertebrates and fishes (Henry and Roberts, 2007; Ross and Quattrini, 2007; Cordes et al., 2008; Rowden et al., 2010; Demopoulos et al., 2014). These characteristics of deep-sea corals make them particularly susceptible to anthropogenic impacts and a priority for conservation efforts.

The degree of connectivity among deep-sea coral populations influences the probability of speciation (Quattrini et al., 2015; Herrera and Shank, 2016) and likely contributes to their ability to respond to natural and anthropogenic stressors. Determining the scale, rate, and directionality of larval dispersal is therefore central to understanding how coral metapopulations are interconnected and the degree of resiliency in the event of a localized disturbance, such as an oil spill (Jones et al., 2007; Almany et al., 2009). Understanding these source-sink dynamics is essential to guide restoration efforts (Lipcius et al., 2008; Puckett and Eggleston, 2016).

Herein, we investigate the spatial patterns of genetic variation and metapopulation connectivity of the octocoral *Paramuricea biscaya* throughout the northern Gulf of Mexico (GoM), using a seascape genomics framework. *Paramuricea biscaya* is one of the most common and abundant corals on hardgrounds on the lower continental slope (between 1200 and 2500 m) in the GoM (Doughty et al., 2014). The 2010 Deepwater Horizon oil spill (DWH) directly impacted populations of this species (White et al., 2012; Fisher et al., 2014; DeLeo et al., 2018) and thus are considered primary targets for restoration (Deepwater Horizon Natural Resource Damage Assessment Trustees, 2016). We use a combination of population seascape genomic analyses, high-resolution ocean circulation modeling, and larval dispersal simulations to quantify the degree of structuring and connectivity among DWH impacted and non-impacted populations. This paper is a companion to the paper by (Liu et al.) that describes the ocean circulation modeling and larval dispersal simulations. Here we test the hypothesis that the genetic diversity of *P. biscaya* is predominantly structured by depth, and to a lesser degree, by distance. We also test the hypothesis that larval dispersal among connected populations is asymmetric due to dominant ocean circulation patterns.

## 2.2 Materials and Methods

### 2.1 Collection of samples

We sampled *Paramuricea biscaya* colonies from six sites in the Northern Gulf of Mexico at depths between 1,371 and 2,400 meters (**Table 1, Figure 1**). The 2010 Deepwater Horizon oil spill directly impacted *P. biscaya* populations at three of these sites in the Mississippi Canyon area (MC294, MC297, and MC344) (White et al., 2012; Fisher et al., 2014). Collections took place during expeditions in 2009 (R/V Ron Brown, ROV Jason II), 2010 (R/V Ron Brown & R/V Atlantis, ROV Jason II & HOV Alvin), 2011 (MSV Holiday Chouest, ROV UHD-34), and 2017 (MSV Ocean Intervention II & MSV Ocean Project, ROV Global Explorer & ROV Comanche). We imaged individual coral colonies before and after removing a small distal branch using hydraulic manipulations mounted on remotely operated vehicles or submarines. We stored samples in insulated containers until the recovery of the vehicles by the surface vessel. Subsamples of each specimen were preserved in liquid nitrogen or 95% ethanol and stored at −80 °C.

**Table 1.** Sampling sites, sample sizes, and environmental characteristics. N_s_ = Sequencing sample size; N_d_ = Dataset sample size after filtering individuals with more than 35% missing data. H_e_ = Nei’s unbiased gene diversity (expected heterozygosity) (Nei, 1978). See the main text for the source of the other environmental parameters.

| Site | $N_s$ | $N_d$ | $H_e$ | Latitude<br>(decimal<br>degrees) | Longitude<br>(decimal<br>degrees) | Depth<br>(m) | Bottom<br>Temperatur<br>e (°C) | Bottom<br>Salinity (S) | Bottom<br>Current Speed<br>(m/s) | Bottom<br>O2<br>(ml/l) | Bottom<br>Potential<br>Density<br>$\sigma_\theta$<br>(kg/m<sup>3</sup>) | Surface<br>Chlorophyll<br>Concentration<br>(mg/m<sup>3</sup>) | Surface<br>Primary<br>Productivity<br>(mg/(m<sup>2</sup> d)) |
| --- | --- | --- | --- | --- | --- | --- | --- | --- | --- | --- | --- | --- | --- |
| DC673 | 37 | 34 | 0.0110 | 28.31174 | -87.30264 | 2254 | 4.39 (0.04) | 35.08 (0.01) | 0.017 (0.006) | 4.21 | 27.835 | 0.22 (0.07) | 326.51 (90.84) |
| GC852 | 34 | 28 | 0.0133 | 27.10997 | -91.16619 | 1407 | 4.26 (0.05) | 35.02 (0.01) | 0.033 (0.007) | 4.36 | 27.792 | 0.15 (0.05) | 258.99 (30.25) |
| KC405 | 31 | 26 | 0.0118 | 26.57086 | -93.48284 | 1679 | 4.25 (0.02) | 35.04 (0.00) | 0.057 (0.011) | 3.87 | 27.814 | 0.15 (0.05) | 269.89 (30.46) |
| MC294 | 18 | 14 | 0.0144 | 28.67225 | -88.47649 | 1371 | 4.78 (0.03) | 34.98 (0.01) | 0.033 (0.004) | 4.29 | 27.706 | 0.59 (0.43) | 551.65<br>(348.93) |
| MC297 | 16 | 13 | 0.0142 | 28.68243 | -88.34401 | 1577 | 4.73 (0.03) | 34.97 (0.01) | 0.046 (0.006) | 4.42 | 27.700 | 0.47 (0.30) | 473.10<br>(221.81) |
| MC344 | 18 | 18 | 0.0128 | 28.63360 | -88.16959 | 1852 | 4.29 (0.01) | 35.02 (0.00) | 0.028 (0.006) | 4.48 | 27.793 | 0.39 (0.24) | 411.74<br>(147.99) |

**Figure 1.**
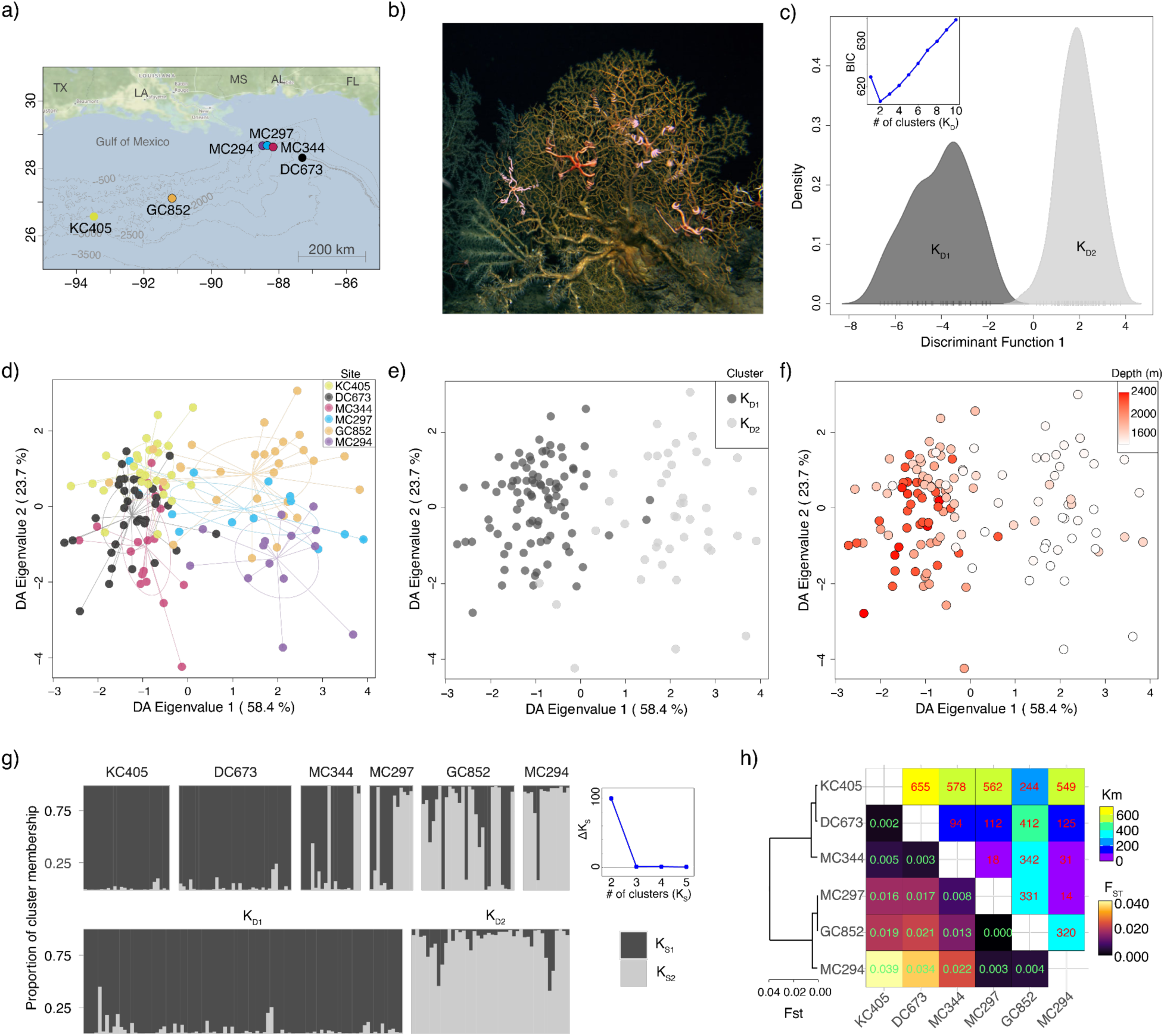
Population genetic structure of *Paramuricea biscaya* in the northern Gulf of Mexico. (**a**) Map showing the study sites in the Gulf of Mexico. (**b**) Image of *Paramuricea biscaya* in its natural habitat in the Gulf of Mexico. (**c**) Density plots of the first discriminant function estimated from DAPC with no sampling location priors. Insert scatter-line plot shows the BIC values for each cluster number (K) tested. K_D1_ and K_D2_ are clusters 1 and 2 identified by DAPC, respectively. (**d**) Scatter plot of the first discriminant analysis (DA) eigenvalues calculated by DAPC with sampling location priors. Individuals are color-coded by sampling site. Colors correspond to sites in panel (a). (**e**) Same scatter plot as in panel (d) but color-coded by DAPC cluster assignment as in panel (c). (**f**) Same scatter plot as in panel (d) but color-coded by individual sampling depth. (**g**) Bayesian population clustering analyses in STRUCTURE. Bars represent individuals grouped by sampling site (top) and DAPC cluster membership (bottom). The color distributions of each bar are proportional to the cluster membership proportions, K_D1_ and K_D2,_ estimated by STRUCTURE. The scatter-line plot on the right shows the ad hoc statistic ΔK, the second-order rate of change of the likelihood function for each cluster number (K) tested. (**h**) Heatmap of pairwise F_ST_ indices of genetic differentiation among sampling sites. **T**he dendrogram on the left was constructed using the Neighbor-Joining algorithm on the F_ST_ values.

### 2.2 Molecular Laboratory Methods

To characterize the genetic diversity of *P. biscaya* individuals, we performed reduced representation DNA sequencing (RAD-seq) (Baird et al., 2008; Reitzel et al., 2013). DNA was purified using the Qiagen DNeasy Blood & Tissue Kit following manufacturers’ protocols. We checked DNA integrity and purity by visual inspection on a 1% agarose gel and a Nanodrop spectrophotometer (Nanodrop Technologies), respectively. DNA concentration was determined and normalized using a Qubit 4.0 fluorometer (Invitrogen). We confirmed species identification through DNA barcoding of the COI mitochondrial gene following the protocols described by (Quattrini et al., 2014) (NCBI GenBank Accession numbers MT795490 to MT795554). Floragenex Inc (Eugene, OR) performed RAD sequencing library preparation utilizing the 6-cutter *PstI* restriction enzyme on quality-checked and concentration-normalized high-molecular-weight DNA. Using the program *PredRAD* (Herrera et al., 2015), we predicted tens of thousands of cleavage sites in coral genomes) with the *PstI* restriction enzyme. Libraries were dual-barcoded and sequenced on an Illumina Hi-Seq 4000 1×100 platform.

### 2.3 Data QC and SNP calling

We de-multiplexed and quality filtered raw sequence RAD-seq reads using the *process_radtags* program in Stacks v2.1 (Catchen et al., 2013) with the following flags: --inline_null, -r, -c, and -q, with default values. We performed read clustering and single nucleotide polymorphism (SNP) calling using the DeNovoGBS (Parra-Salazar et al., 2021) module of the software package NGSEP v4.0.1 (Tello et al., 2019). This software is more computationally efficient and has comparable or better accuracy than programs like Stacks or pyRAD (Eaton, 2014) for de novo analysis of genotype-by-sequencing data (Parra-Salazar et al., 2021). We assumed a heterozygosity rate of 1.5% (-h 0.015) as calculated from the short read genome-wide data of the sister species *Paramuricea* B3 using the software GenomeScope v2.0 (Vurture et al., 2017) from the National Center for Biotechnology Information (NCBI) Sequence Read Archive (SRA) under BioProject number PRJNA574146 (Vohsen et al., 2020).

### 2.4 SNP and Individual Filtering

Single nucleotide polymorphisms were filtered using vcftools v0.1.16 (Danecek et al., 2011) to exclude SNP loci that: 1) had more than 30% missing data, 2) a mean depth of coverage smaller than 10x or greater than 1000x, 3) a minor allele frequency smaller than 0.01, and 4) had more than two alleles. The resulting dataset, containing 12,948 SNPs and 154 individuals, is hereafter referred to as the *all_snp* dataset. *BayeScan* v.2.01 (Foll and Gaggiotti, 2008) was used to identify SNP potentially under positive selection (-n 5,000 -burn 50,000 -pr_odds 10,000, Qval<0.05).

The *all_snp* dataset was imported into the R v4.0.3 statistical environment (Team and Others, 2013) for further filtering. We excluded individuals if they had missing data in more than 35% of the SNP loci or identified as clones by *clonecorrect* function from the R package *poppr* v2.8.6 (Kamvar et al., 2014). We excluded SNP loci if their observed heterozygosity was greater than 0.5, as estimated with *hierfstat* v0.5 (Goudet, 2005), or if their allelic frequencies were not in Hardy-Weinberg equilibrium, as estimated with *pegas* v0.14 (Paradis, 2010) (B=1000, p<0.01). We randomly retained one SNP per RAD locus to reduce the risk of violating the assumption of independence among SNP. Finally, 10 SNPs in RAD loci identified as potentially under positive selection by *BayeScan* were excluded (**Supplementary figure S1**). This dataset, containing 4,248 unlinked neutral SNPs across 133 individuals, is hereafter referred to as the *neutral* dataset.

### 2.5 Genetic connectivity

To measure the genetic connectivity among sampling sites, we estimated migration rates (*m*), defined as the proportion of immigrant individuals in the last two generations, using BAYESASS v3.0.4.2 (Wilson and Rannala, 2003). Twelve independent runs with different random seeds were performed using the *neutral* dataset. We ran each analysis for 100 million Markov chain Monte Carlo (MCMC) iterations, with 50 million burn-in iterations and one thousand iterations sampling frequency. Mixing parameters (-m0.35 -a0.9 -f0.09) were optimized to ensure adequate mixing (acceptance rates between 20 and 60%). MCMC trace files were examined in the program Tracer v1.7.1 (Rambaut et al., 2018) to evaluate convergence and consistency of estimates among runs. We calculated point estimates of *m* as the median of the posterior distribution and their uncertainty as 95% High Posterior Density (HPD) intervals.

### 2.6 Potential connectivity

To identify dispersal mechanisms that could explain genetic connectivity estimates, we compared our results with the potential connectivity estimates (probability of connectivity through larval dispersal among sampling sites) by Liu et al. (submitted). Briefly, Liu et al. (submitted) simulated the dispersal trajectories of neutrally-buoyant Lagrangian particles in an implementation of a high-resolution three-dimensional Coastal and Regional Ocean COmmunity hydrodynamic model (CROCO) (for full details, see (Liu et al.)). The model encompassed the area between 98o-82o W and 24o-31o N and had a horizontal grid resolution of approximately 1 km and 50 vertical sigma (density) layers. 4489 Lagrangian particles were deployed uniformly at the seafloor in 0.05 × 0.05º areas centered at the location of the sampling sites. The particles were tracked offline using the Lagrangian tool Ichthyop (Lett et al., 2008) and recorded hourly. Horizontal connectivity through larval dispersal among sampling sites (*l*_*h*_) was defined as the average proportion of neutrality-buoyant Lagrangian particles released at a source site (*i*) area that passed over another site (*j*) area (sink) after 56 days (computational constraints limited the length of the tracking) starting from January 25th, April 25th, July 24th, and November 1st, 2015. The pelagic larval duration (PLD) for Paramuricea biscaya is unknown, but Hilario et al. ((Hilario et al., 2015) found that a PLD between 35 and 69 days seems representative of 50% to 75% of deep-sea species. The definition of vertical connectivity (*l*_*v*_) is the same as horizontal connectivity, except that a particle also has to pass within 50 meters of the sink site’s seafloor depth. Liu et al (submitted) also evaluated longer PLDs by extending the Lagrangian tracking starting November 1st to 148 days and with additional Eulerian dye releases followed for 120 days. The dye release indicated that although Lagrangian particles cover a smaller area than the Eulerian dye, they capture the same main dispersal features and do not predict substantially different connectivity patterns. No other biological parameters such as larval growth, mortality, settlement, and swimming because they are unknown for the study species.

### 2.7 Population genetic structure

To determine the patterns of genetic structuring of the sampled *P. biscaya* corals, we performed a discriminant analysis of principal components (DAPC) on the *neutral* dataset using the R package *adegenet* v2.1.3 (Jombart, 2008). DAPC was performed with and without sampling locations as priors after estimating the optimal number of principal components with the function *optim*.*a*.*score*. For the DAPC with no priors, we applied the Bayesian Information Criterion to choose the optimal number of clusters (K) that explain the genetic variability in the dataset using the function *find*.*clusters*.

We also inferred population structuring patterns (as historical lineages) with the *neutral* dataset by maximizing the posterior probability of the genotypic data, given a set number of clusters (K). This method is known as Bayesian population clustering and is implemented in the program *structure* v2.3.4 (Pritchard et al., 2000). We used the admixture model with uncorrelated allele frequencies. The MCMC was run for 1.1×10^6^ repetitions (burn-in period 1×10^5^). We evaluated values for K from 1 to 6 (10 replicates each). We selected the optimal value of K using the program *StructureHarvester* v0.6.92 (Earl and vonHoldt, 2012) according to the *ad hoc* ΔK statistic (Evanno et al., 2005), which is the second-order rate of change of the likelihood function. We visualized *structure* results using the program R package *starmie (Tonkin-Hill and Lee, 2016)*.

We performed a hierarchical Analysis of Molecular Variance (AMOVA) (Excoffier et al., 1992) with the *neutral* dataset to calculate F-statistics and test for differentiation at the individual, site, and genetic cluster levels. The AMOVA, performed in *genodive* v3.04 (Meirmans, 2020), assumed an infinite-alleles model. We calculated pairwise F_ST_ (Weir and Cockerham, 1984) differentiation statistics among sampling sites with the R package *assigner* v.0.5.8 (Gosselin).

### 2.8 Redundancy Analyses

To quantify environmental variables’ significance and relative importance in shaping genetic diversity in *P. biscaya*, we used a series of redundancy analyses (RDA) in the R package *vegan* v2.5 (Oksanen et al., 2007). RDA has two steps. First, a multiple linear regression between genetic (response) and environmental (explanatory) data matrices produces a matrix of fitted values. Second, a principal components analysis (PCA) of the fitted values. The PCA axes are linear combinations of the explanatory variables (Legendre and Legendre, 2012).

We performed site-level RDA (Legendre and Legendre, 2012) on sites’ allelic frequencies and geographical distances. We first transformed geographical distances as in-water distances using the *lc*.*dist* function of the R package *marmap* v1.0.4 (Pante and Simon-Bouhet, 2013), and later represented as distance-based Moran’s eigenvector maps (dbMEM) (Dray et al., 2006) using the R package *adespatial* (Dray et al., 2018).

We performed individual-level distance-based RDA (dbRDA) (McArdle and Anderson, 2001) with the matrix of genetic distances calculated from the *neutral* dataset and a matrix of environmental variables. Missing genotypes in each individual were first imputed by assigning the most common genotype for each locus at the collection site. Environmental variables included: depth, latitude, longitude, bottom temperature, salinity, bottom current speed, bottom oxygen concentration, bottom seawater potential density (σ_θ_), surface chlorophyll concentration, and surface primary productivity.

Average monthly bottom temperature, salinity, current speed values between 2011-2018, as extracted by Goyert et al. ((Goyert et al.) from the HYbrid Coordinate Ocean Model (HYCOM) for the Gulf of Mexico, were summarized as mean and standard deviation grids with a 4 km resolution. Bottom oxygen concentration values are annual means gridded from the World Ocean Database by Goyert et al. ((Goyert et al.) at a 370 m resolution. We calculated bottom seawater potential density (σ_θ_) values using the R package *oce* v1.2 (Kelley, 2018). We obtained average monthly surface chlorophyll concentration and primary productivity values between 2011-2018 from the E.U. Copernicus Marine Service Information, Copernicus Globcolour ocean products grids OCEANCOLOUR_GLO_CHL_L4_REP_OBSERVATIONS_009_082 and OCEANCOLOUR_GLO_OPTICS_L4_REP_OBSERVATIxONS_009_081, and summarized as mean and standard deviation grids with a 4 km resolution. Individual parameter values were extracted from these grids using the latitude and longitude of each sampled coral.

To avoid problems with highly correlated environmental variables (Dormann et al., 2013), we performed a pairwise correlation test and removed variables with a correlation coefficient |r| > 0.7 and a p-value < 0.05. We retained the most seemingly ecologically relevant variable when two or more variables were correlated. We evaluated the explanatory importance of each environmental variable using forward selection and analysis of variance (ANOVA) after 10,000 permutations (**α** = 0.05) using the *ordistep* function in *vegan*. Retained environmental variables were included in the dbRDA using the *dbrda* function in *vegan*. We performed a variance partitioning analysis with the function *varpart* and tested its significance through global and marginal ANOVAs (1,000 permutations, **α** = 0.01).

## 3. Results

### 3.1 Population Connectivity

Migration rates (*m*) among sites estimated from genetic data using BAYESASS were overall low (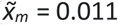, s_m_ = 0.065), with a few exceptions. Approximately 20% of individuals at the sites MC344 (depth 1852 m) and KC405 (1679 m) likely immigrated from the De Soto Canyon area (site DC673, depth 2254 m), within the last one or two generations (**Figure 2a, Supplementary Figure S2**; *m*_*DC673-MC344*_ = 0.210, 95% HPD_*DC673-MC344*_ = [0.145,0.269]; *m*_*DC673-KC405*_ = 0.181, 95% HPD_*DC673-KC405*_ = [0.123,0.236]). The potential contribution of immigrants from DC673 to site GC852 (depth 1407 m), in the Green Canyon area, and sites MC294 (depth 1407 m)and MC297 in the Mississippi Canyon area was smaller but still substantial, ranging between 3 and 10% (**Figure 2a, Supplementary Figure S2**; *m*_*DC673-GC852*_ = 0.097, 95% HPD_*DC673-GC852*_ = [0.050,0.149]; *m*_*DC673-MC294*_ = 0.051, 95% HPD_*DC673-MC294*_ = [0.000,0.076]; *m*_*DC673-MC297*_ = 0.085, 95% HPD_*DC673-MC297*_ = [0.028,0.153]). These analyses also indicate that GC852 may also be an important source of immigrants to the Mississippi Canyon area. The potential contribution of immigrants from GC852 to sites MC344, MC297, and MC294 ranges between 5 and 24% (**Figure 2a, Supplementary Figure S2**; *m*_*GC852-MC344*_ = 0.052, 95% HPD_*DC673-GC852*_ = [0.012,0.105]; *m*_*GC852-MC294*_ = 0.236, 95% HPD_*DC673-MC294*_ = [0.167,0.295]); *m*_*DC673-MC297*_ = 0.139, 95% HPD_*DC673-MC297*_ = [0.071,0.212]).

**Figure 2.**
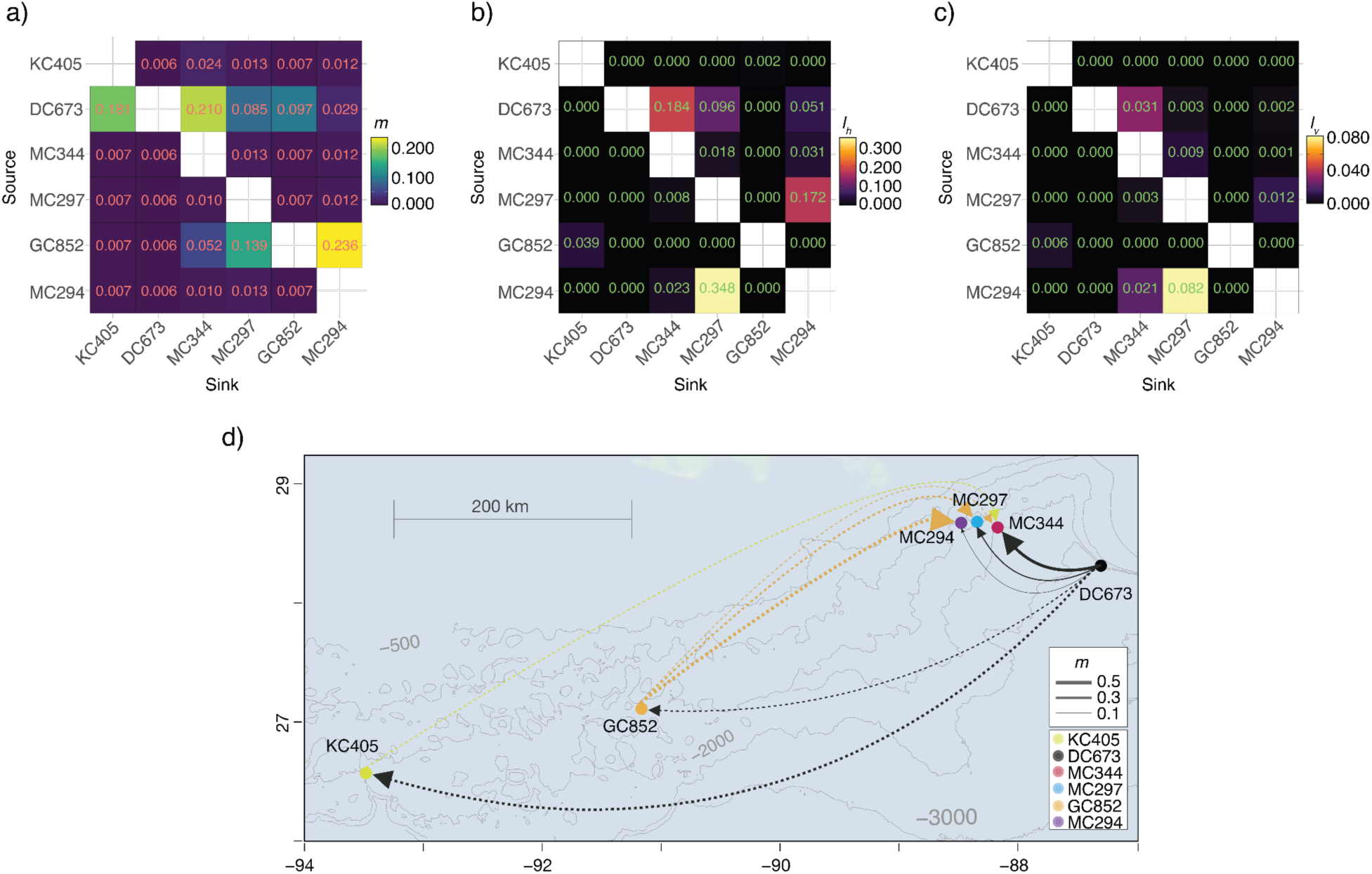
Directional population connectivity *Paramuricea biscaya* in the northern Gulf of Mexico. a) Migration rates (*m*) inferred from observed genetic data using BAYESASS. b) Horizontal connectivity probabilities (*l*_*h*_) integrated over all available periods (56 days in February, April, and August releases, and 148 days for November) calculated from larval dispersal simulations. c) 3D connectivity probabilities (*l*_*v*_) (including both horizontal and vertical components) calculated from larval dispersal simulations. Rows in each matrix indicate source sites, and columns indicate sink sites. Figures b) and c) are modified from Liu et al (submitted). d) Map depicting population connectivity patterns among study sites. Dots indicate sites. A line connecting two dots indicates an observed (genetic) or predicted (model) connection >0. Solid lines indicate connections supported by both genetic and model data. Dashed lines indicate connections supported only by genetic data. Arrowheads indicate the direction of the connection. Line thickness is proportional to the strength of the connection (measured as migration rate *m*). Line and dot colors indicate site identities and correspond to those in Figures 1a and 1d.

Horizontal connectivity probabilities (*l*_*h*_) calculated from larval dispersal simulations recovered a remarkable congruence with the estimated migration rates (*m)* concerning the role of the De Soto Canyon area DC673 as a source of larvae for the Mississippi Canyon sites (**Figure 2b and 2d**, *l*_*h DC673-MC344*_ = 0.184, ; *l*_*h DC673-MCC294*_ = 0.051; *l*_*h DC673-MC297*_ = 0.096), but not for the Green Canyon (GC852) or the Keathley Canyon (KC405) areas. The congruence is only maintained in the 3D connectivity probability (*l*_*v*_) for MC344 (**Figure 2c-d**, *l*_*v DC673-MC344*_ = 0.031). The larval dispersal simulations also predict bi-directional connectivity between MC294 and MC297 (*l*_*h MC294-MC297*_ = 0.348, ; *l*_*h MC297-MCC294*_ = 0.172; *l*_*v MC294-MC297*_ = 0.082; *l*_*v MC297-MCC294*_ = 0.012) but the estimated migration rates between these sites are low (**Figure 2a, Supplementary Figure S2;** *m*_*MC294-MC297*_ = 0.013, 95% HPD_*MC294-MC297*_ = [0.000,0.051]; *m*_*MC297-MCC294*_ = 0.012, 95% HPD_*MC297-MCC294*_ = [0.000,0.049]).

### 3.2 Population Genetic Structure

DAPC analysis with no location priors indicated that there is metapopulation substructuring within *P. biscaya’s* sampled range. The variability in the genetic data was explained by two clusters K_D1_ (n = 89) and K_D2_ (n = 44) (optimal K=2, BIC=616.8, 18 retained PCs (**Figure 1c**). Each individual was considered a member of the group with the highest probability. The first cluster is mainly composed of individuals collected at sites DC673 (100% of sampled individuals belong to K_D1_), KC405 (100% K_D1_), and MC344 (83% K_D1_), while the second cluster is mainly composed of individuals collected at sites MC294 (93% of sampled individuals belong to K_D2_), MC297 (54% K_D2_) and GC852 (75% K_D2_) (**Figures 1c-e**). The first discriminant axis, calculated by DAPC analysis with location priors, explained 58.4% of the variance and primarily reflected the differentiation between the two inferred clusters K_D1_ and K_D2_. This differentiation seemed to be associated with depth as samples assigned to K_D1_ were on average found at deeper locations (mean depth at which K_D1_ individuals were sampled: 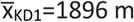, standard deviation: s_KD1_= 308 m) than individuals assigned to K_D2_ (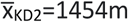, s_KD2_=130m) (**Figure 1f**).

The STRUCTURE analyses of Bayesian population clustering confirmed the presence of two ancestry clusters, K_S1_ (n = 91) and K_S2_ (n = 42), that largely corresponded to the clusters identified by the DAPC K_D1_ and K_D2_, respectively (we considered each individual a member of the group for which it had the highest membership probability *Q*). To maintain consistency with other studies in corals (Carlon and Lippé, 2011; Serrano et al., 2016), we defined an admixed individual as having a *Q* > 0.1 for both clusters. These analyses indicate that, overall, 15% of individuals have an admixed ancestry (**Figures 1g**), but proportionally there are more admixed individuals assigned to K_S2_ (24% of individuals) than to K_S1_ (11%). Ancestry cluster K_S1_ is dominant in DC673 (mean probability of membership a that site: 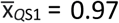, KC405 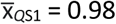 and MC344 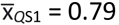, whereas K_S2_ is dominant in MC294 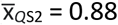, GC852 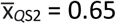, and MC297 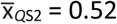.

Pairwise F_ST_ (Weir and Cockerham, 1984) statistics among sites were consistent with the DAPC and STRUCTURE results showing greatest differentiation among sites with a majority of individuals assigned to different K clusters, and lowest among sites with a majority of individuals assigned to the same K cluster (**Figures 1h**). The AMOVA analysis indicated that 11.4% of the observed genetic variation could be attributed to differences among individuals (F_IS_ = 0.117, p = 0.001), 0.3% to differences among sites (F_SC_ = 0.003, p = 0.001) and 1.8% to differences between DAPC clusters K_D1_ and K_D2_ (F_CT_ = 0.018, p = 0.001).

### 3.3 Redundancy Analyses

Site-level RDA failed to detect a significant correlation (**α** = 0.05) between geographic distance (as dbMEM eigenvectors) and genetic differentiation, thus rejecting the hypothesis of isolation by distance.

Of the environmental variables, we excluded bottom temperature and salinity from the individual-level dbRDA as the ranges of these parameters across the study sites were too small to be biologically important (4.21 to 4.84 °C, and 34.95 and 35.10 PSU). Mean surface primary productivity (retained) was significantly correlated with its standard deviation and latitude and mean and standard deviation of surface chlorophyll concentration. Depth (retained) was significantly correlated with bottom seawater potential density (σ_θ_). Mean bottom oxygen concentration (retained), longitude (retained), and mean bottom current speed (retained) were not significantly correlated with any other environmental variable.

We incorporated depth, mean bottom dissolved oxygen concentration, mean surface primary productivity, longitude, and mean bottom current speed into an initial dbRDA model as these were the only significant independent variables identified by forward selection (ANOVA, p-values < 0.05). These variables significantly contributed to the model, except for mean bottom current speed (ANOVA, p-value > 0.01), which was subsequently excluded.

Globally, the percentage of the genetic variation explained by environmental variables was 7.37% (**Table 2**). Depth (collinear with bottom seawater potential density (σ_θ_)) had the largest effect (explaining 3.8% of the variance), followed by mean bottom oxygen concentration (2.0%), mean surface primary productivity (collinear with six other variables, see above) (1.8%), and longitude (1.1%). The combined effect of depth and density is evident in the dbRDA plots (**Figure 3**). dbRDA axes are linear combinations of the environmental variables. dbRDA axis 1, which explains 52.5% of the variation, broadly splits individuals belonging to different DAPC clusters. This differentiation is primarily driven by depth as indicated by the environmental variables vectors and suggested in **Figure 1e-f**.

**Table 2.** Environmental variables tested in the dbRDA.

| Variable | Df | Variance Partition |  | ANOVA |  |  |
| --- | --- | --- | --- | --- | --- | --- |
|  |  | R<sup>2</sup> | Adjusted R<sup>2</sup> | Variance | F | Pr(>F) |
| Depth | 1 | 0.0453 | 0.0380 | 73.60 | 2.6038 | 0.0010 |
| Bottom Dissolved O<sub>2</sub> Concentration | 1 | 0.0274 | 0.0200 | 41.40 | 1.4627 | 0.0050 |
| Mean Surface Primary Productivity | 1 | 0.0252 | 0.0178 | 43.40 | 1.5337 | 0.0010 |
| Longitude | 1 | 0.0186 | 0.0111 | 51.00 | 1.8033 | 0.0010 |
| All (global) | 4 | 0.1017 | 0.0737 | 273.10 | 2.4138 | 0.0010 |

**Figure 3.**
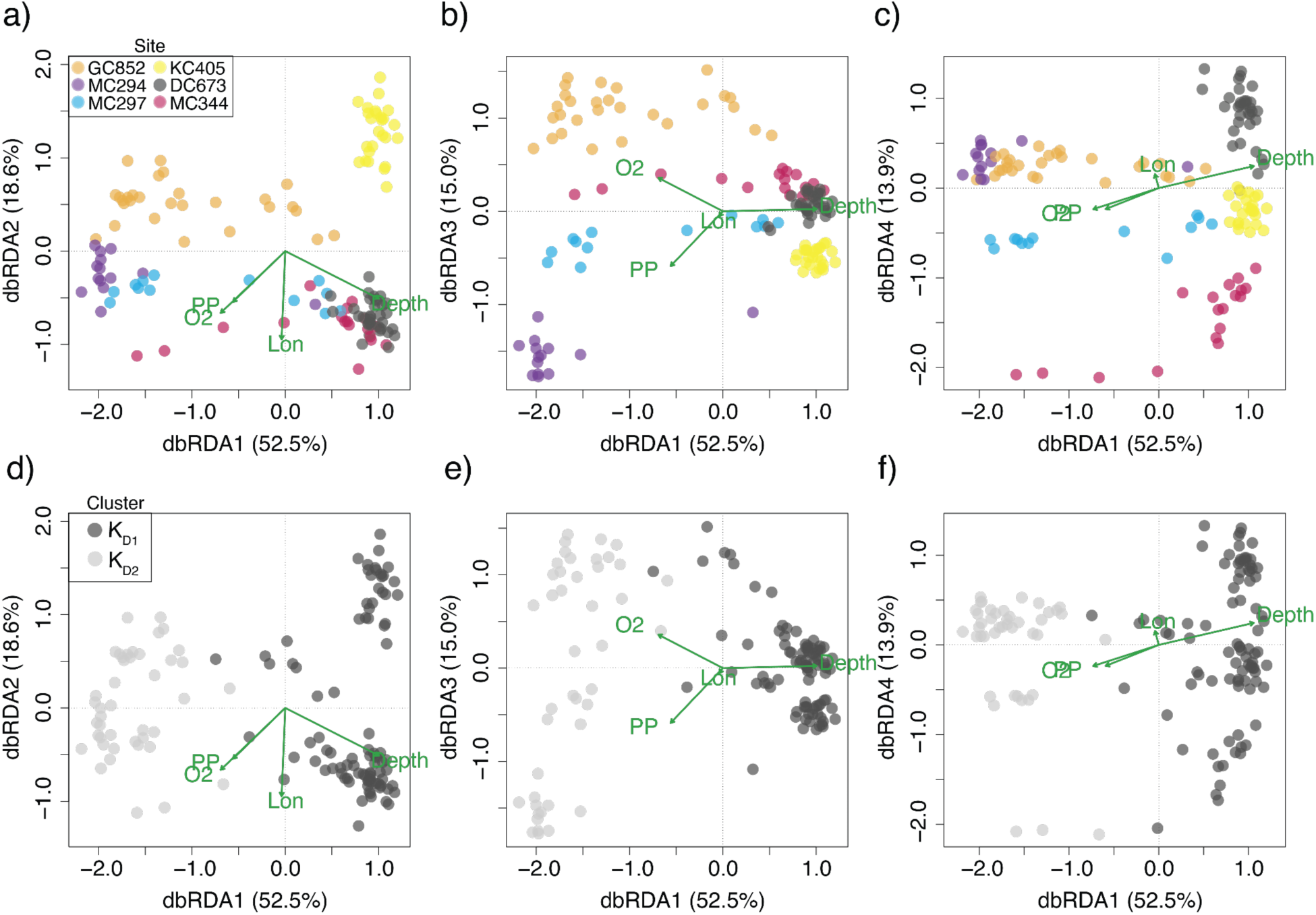
Diplots of distance-based Redundancy Analysis (rbRDA) of environmental (explanatory) and genetic (response) variables. Green vectors represent environmental variables: depth, bottom dissolved oxygen concentration (O_2_), surface primary productivity (PP), and longitude (Lon). The length of each vector is proportional to its contribution to each axis. rbRDA axes are linear combinations of the environmental variables. Dots represent individual corals. Individuals are color-coded by a-c) sampling site and d-f) DAPC cluster assignment.

## 4. Discussion

### 4.1 Population Connectivity: Scale, Rate, and Directionality

Larval dispersal simulations in the study area show a prevailing westward pathway of dispersal along isobaths in the 1,000-2,000 m range in all seasons (Liu et al.). Long-distance dispersal (more than 100 km) driven by strong deep recirculation currents (Bracco et al., 2016) may occur for larvae originating in the DeSoto Canyon area (DC673) (Liu et al.). These larvae can reach the Mississippi Canyon area in less than two months (Liu et al.), thus explaining the source-sink dynamics identified between these sites by migration rate estimates (*m)* from genetic data (**Figure 2**). These source-sink dynamics are highly depth-dependent. Our estimates suggest that 15 to 27% of individuals at MC344 (1852 m) likely immigrated from the De Soto Canyon area (DC673, 2254 m) within the last one or two generations. For MC297 (1577 m), 3 to 15% are likely immigrants from DC673, and less than 8% for MC294 (1371 m). The limiting effect of depth on vertical connectivity is most striking within the Mississippi Canyon (**Figure 2**), where we found no evidence of substantial gene flow among sites. The limited amount of vertical diapycnal mixing possible over the short horizontal distances that separate them (tens of kilometers, see the following section for further discussion on the role of depth) may explain the limited 3D connectivity among these sites (Bracco et al. 2019; Liu et al.).

Our analyses indicate that the population of *P. biscaya* at DC673 should be a conservation priority to restore the impacted populations at MC344 and MC297. Additional sampling and modeling throughout *P. biscaya’s* depth range (1,000 to 2,600 meters) in the DeSoto Canyon and West Florida Escarpment are necessary to fully understand the role of this region as a source of larvae for DWH impacted populations in the Mississippi Canyon and identify other sites in need of protection.

Larval dispersal models predict that larvae originating from the Keathley Canyon area (KC405) can disperse the furthest (maximum horizontal distances 154km and 426km after 56 days and 148 days, respectively (Liu et al.)). The highly variable currents that characterize this area can explain this potential for long-distance dispersal (Liu et al.). However, these models fail to predict the degree of direct genetic connectivity estimated between KC405 and DC673 (**Figure 2**). Similarly, the relative importance of the Green Canyon site GC852 as a source of larvae to the Mississippi Canyon area, indicated by the migration rate estimates, is not consistent with the connectivity probabilities estimated by the numerical larval simulations (Liu et al.). The dispersal distances for larvae out of GC852 do not seem to exceed 100 km after 56 days (400 km after 148 days) (Liu et al.). Thus no direct connectivity is predicted between the Green Canyon and Mississippi canyon sites separated by more than 300 km.

The patterns of genetic connectivity between KC405 and DC673, and GC852 and the Mississippi Canyon sites cannot be explained by larval dispersal models unless intermediate populations that act as stepping stones are included in the simulations (Liu et al.) when the role of interannual variability is accounted for by using the advection pathways predicted by HYCOM data. Additional targeted exploration and sampling, informed by habitat suitability (Georgian et al., 2020) and dispersal models (Liu et al.), are necessary to test this connectivity hypothesis and clarify the role of western populations in the restoration of DWH impacted populations.

### 4.2 Metapopulation Structuring by Depth

All of our analyses support the existence of two clusters or ‘stocks’ of *Paramuricea biscaya* in our samples, both of which were impacted by the DWH oil spill. Previous studies sequenced mitochondrial DNA of *P. biscaya* (*mtCOI+igr+MutS*) and recovered three haplotypes of *P. biscaya* (B1, B1a, B2) in the northern Gulf of Mexico (Doughty et al., 2014; Quattrini et al., 2014; Radice et al., 2016). We found that mitochondrial haplotypes bear no direct correspondence with the genomic clusters (Supplemental Figure S3). Mitochondrial markers are well known for lacking sufficient variability at low taxonomic levels in octocorals and are subject to incomplete lineage sorting (Pante et al., 2015; Herrera and Shank, 2016; Quattrini et al., 2019). We suggest that mitochondrial DNA barcoding data should not be used to resolve differences at the population level, especially in the context of management and restoration, and in many cases at the species level, as it could lead to incorrect interpretations and inadequate policy decisions.

Geographic distance is not a significant variable structuring the genetic diversity of *P. biscaya* within the GoM. Despite only being separated by tens of kilometers, the populations in the Mississippi Canyon impacted by the DWH oil spill (MC294, MC297, and MC344) have distinct genetic compositions (**Figure 1**). The population’s genetic composition at MC344 is most similar to those found at the DeSoto Canyon (DC673) and Keathley Canyon (KC405), hundreds of kilometers away. Consistent with results from previous studies of deep-sea populations (Taylor and Roterman, 2017), depth is a critical variable structuring the genetic diversity of *P. biscaya*. MC344 is the deepest of the three sites at the Mississippi Canyon (MC294: 1371 m, MC297: 1577 m, and MC344: 1852 m), and its population is mainly composed of individuals whose ancestry is predominantly from the first cluster K_D1_ (83%; the other deep sites DC673 [2254 m] and KC405 [1679 m] are also almost entirely made up of individuals with K_D1_ ancestry). MC294, the shallowest, has a population whose ancestry is mainly from the second cluster, K_D2_ (93%). MC297 sits at an intermediate depth and has a population of mixed ancestry, split roughly in half.

Seascape genomic analyses provide statistical support for the role of depth. The combined effect of depth and bottom seawater potential density (σ_θ_) contributes the most towards explaining the genetic variability in *P. biscaya* among the environmental variables explored in this study (**Figure 3, Table2**). Among the environmental variables known to show collinearity with depth, hydrostatic pressure may be the most biologically important for *Paramuricea biscaya*. Hydrostatic pressure increases linearly with depth (at a rate of roughly 1 atmosphere every 10 meters). Other variables, such as dissolved oxygen concentration, pH, temperature, and salinity, do not vary sufficiently within the depth and geographical range of the examined populations in the study area to exert any significant adaptive pressure that could drive diversification. Several studies have suggested that pressure can be a significant selective force in the deep sea, often driving the evolution of pressure-adapted enzymes and other biomolecules (Somero, 1992; Lan et al., 2017, 2018; Gaither et al., 2018; Lemaire et al., 2018; Gan et al., 2020; Weber et al., 2020).

Another potentially important variable known to be collinear with depth is the flux of particulate organic matter from the surface ocean to the seafloor (POC flux). POC is the primary food source for most deep-sea organisms, and it is known to structure biodiversity patterns on the benthos (Woolley et al., 2016). POC flux decreases exponentially with depth (Martin et al., 1987; Mouw et al., 2016) and could therefore be a significant selective force (Quattrini et al., 2017) and a major driver of biodiversity patterns in the deep sea. However, POC accumulates on the seafloor, where it can be resuspended through the interaction of bottom currents and complex topography (Wilson et al., 2015; Amaro et al., 2016). The role of POC resuspension is uncertain given the diversity of habitats where *P. biscaya* is found, from carbonate outcrops (e.g., GC852 and MC sites) to near-vertical walls (DC673, KC405). Furthermore, episodical delivery episodes of POC are challenging to incorporate in models but are likely biologically important (Smith et al., 2018). In-situ measurements would be needed to quantify differences in food delivery at these sites. Although chlorophyll-a concentrations, sea surface temperature, and photosynthetically active radiation are used to model net primary productivity (Behrenfeld and Falkowski, 1997) and POC fluxes (Pace et al., 1987), their relationship is not always predictable. In the northern Gulf of Mexico, confounding factors such as planktonic community composition can cause discrepancies between modeled and *in situ* POC flux measurements (Biggs et al., 2008; Maiti et al., 2016).

Bottom seawater potential density (σ_θ_) could play an important role if larvae behave as neutrally buoyant particles dispersing along isopycnals as suggested for other deep-sea corals and sponge species (Dullo W et al., 2008; Kenchington et al., 2017; Bracco et al., 2019; Roberts et al., 2021). Larval dispersal along narrow density envelopes associated with water mass structuring may serve as a mechanism for increasing reproductive success and gene flow among deep-sea metapopulations of species with neutrally buoyant larvae while simultaneously facilitating pre-zygotic isolation by limiting dispersal across depth (Miller et al., 2011). The results from our potential connectivity analyses further suggest that the ocean circulation, and specifically the limited diapycnal mixing, may prevent neutrally-buoyant larvae from spreading across depth ranges.

Remarkably, all sampled sites, except for DC673, have a proportion of individuals with admixed ancestry, suggestive of successful crosses beyond F1 or F2 generations among clusters (**Figure 1**). This could be indicative of an absence of post-zygotic isolation barriers between clusters. There are two possible explanations for this pattern: incipient sympatric speciation or secondary contact. Incipient sympatric ecological speciation through niche specialization is a possible driver of the observed pattern of population structuring (González et al., 2018). Due to the relative environmental stability at the depth range of *P. biscaya*, it is plausible that specialization to pressure and food gradients would occur over the species’ depth range and be reinforced by density-driven dispersal limitation. Alternatively, secondary contact could occur by recent colonization of the GoM by a *P. biscaya* lineage from the Caribbean Sea or the Atlantic Ocean. Differentiating between the two possibilities would require a combination of demographic modeling and additional sampling throughout the range of *P. biscaya*, i.e. not limited to the GoM.

The presence of these two genetic stocks should be taken into consideration for restoration activities that involve propagation in nurseries and transplantation (Baums et al. 2019). The possibility that the stocks are partially reproductively isolated and depth-adapted suggests that receiving populations would benefit most from transplants from populations to which they are already genetically connected.

## 5. Conclusions

In this study, we found support from population genomic analyses and larval dispersal modeling for the hypothesis that the *P. biscaya* metapopulation in the northern GoM is predominantly structured by depth, and to a lesser degree, by distance. Further, both lines of evidence (genetic and modeling) support the hypothesis that larval dispersal among connected populations is asymmetric due to dominant ocean circulation patterns. Utilizing a seascape genomic approach brought a more holistic understanding of the population connectivity of this species than either population genetics or modeling could on its own. There are likely intermediate unsampled populations that serve as stepping stones for dispersal. These may explain some of the observed genetic connectivity that could not be explained as direct dispersal by larval simulations. Although dispersal and connectivity patterns of organisms are highly species-dependent, the integrative framework of this study provides valuable insights to understand the connectivity of deep-sea metacommunities broadly (Mullineaux et al., 2018).

This study further illustrates that management of marine protected areas (MPAs) should incorporate connectivity networks and depth-dependent processes throughout the water column. Doing so could help preserve genetic diversity and increase species resilience to extreme climate events and anthropogenic impacts. We suggest that the DeSoto Canyon area, and possibly the West Florida Escarpment, critically act as sources of larvae that may repopulate areas impacted by the 2010 Deepwater Horizon oil spill in the Mississippi Canyon. Active management of these source sites is essential to the success of restoration efforts.

## Author Contributions

SH, AMQ and AB, designed the research. SH led the field work and project management. SH, MPG, DW, AB, and GL performed the research. SH, MPG, AB and GL analyzed data. SH wrote the paper with contributions from MPG, AB, GL and AMQ.

## Acknowledgments

We thank Chuck Fisher, Erik Cordes, Illiana Baums for leading supplemental field efforts and providing access to samples and dive time. We thank Matthew Potti and Michael Coyne for providing access to HYCOM and oxygen grids. We would also like to thank Sam Vohsen, Alexis Weinnig, Janessy Frometa, Fanny Girard, Amanda Glazier, Amanda Demopoulos, and the crew of expeditions We would also like to thank Alondra Maldonado for her help with DNA purifications, and Dan Fornari and Peter Etnoyer for assisting with field logistics. We thank Cathy McFadden, Cheryl Morrison, and Frank Parker for project support.

## Data accessibility

Coral samples are housed in the Herrera Lab at Lehigh University. Raw RAD-seq sequence data is available at the NCBI SRA database under BioProject number PRJNA766840. COI barcodes have been submitted to NCBI: MT795490 to MT795554. The SNP datasets, environmental matrix and individual sampling information have been deposited at FigShare 10.6084/m9.figshare.16692229.

## Supplementary Material

**Figure S1.**
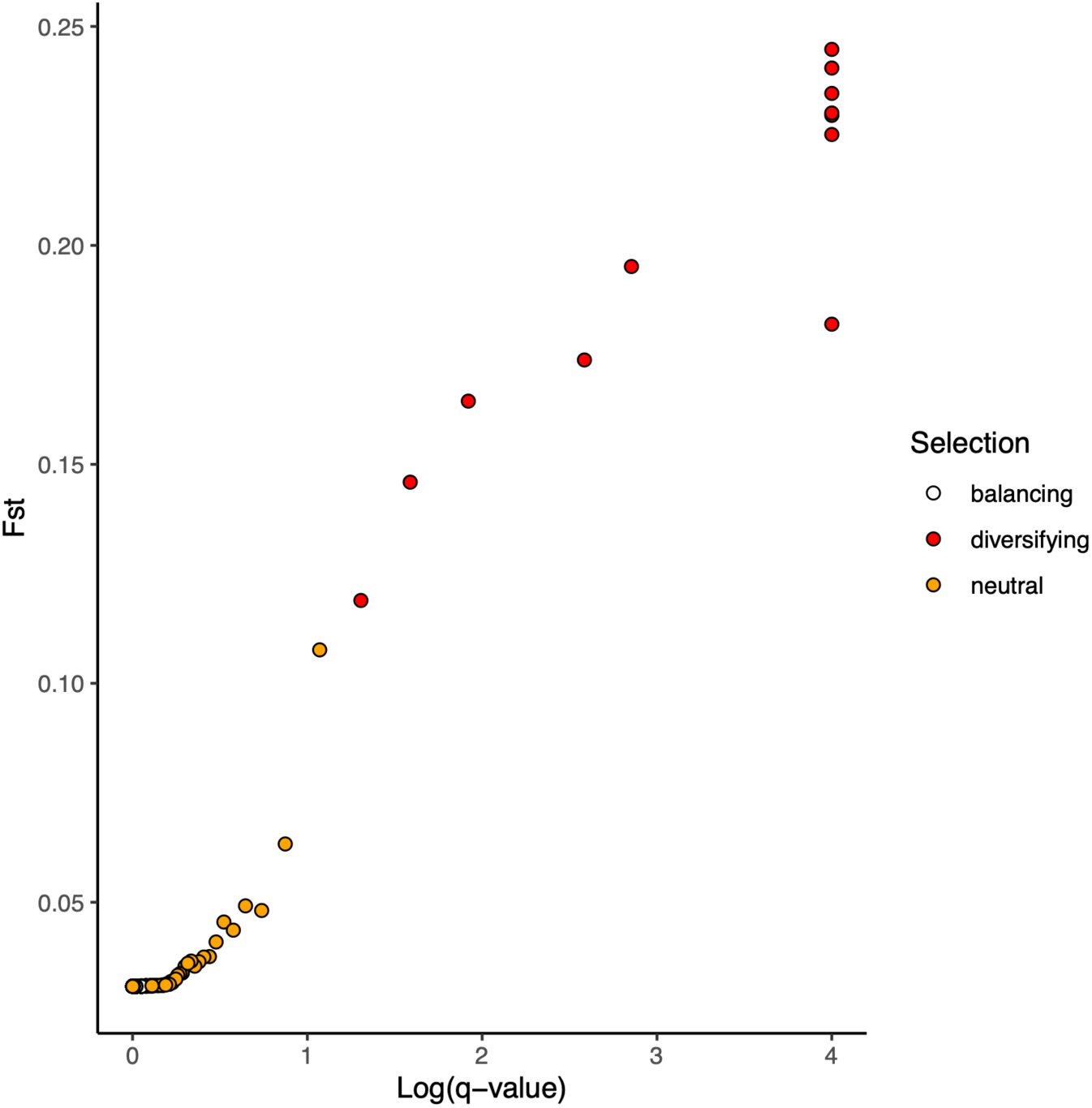
Bayescan plot showing outlier loci under diversifying selection

**Figure S2.**
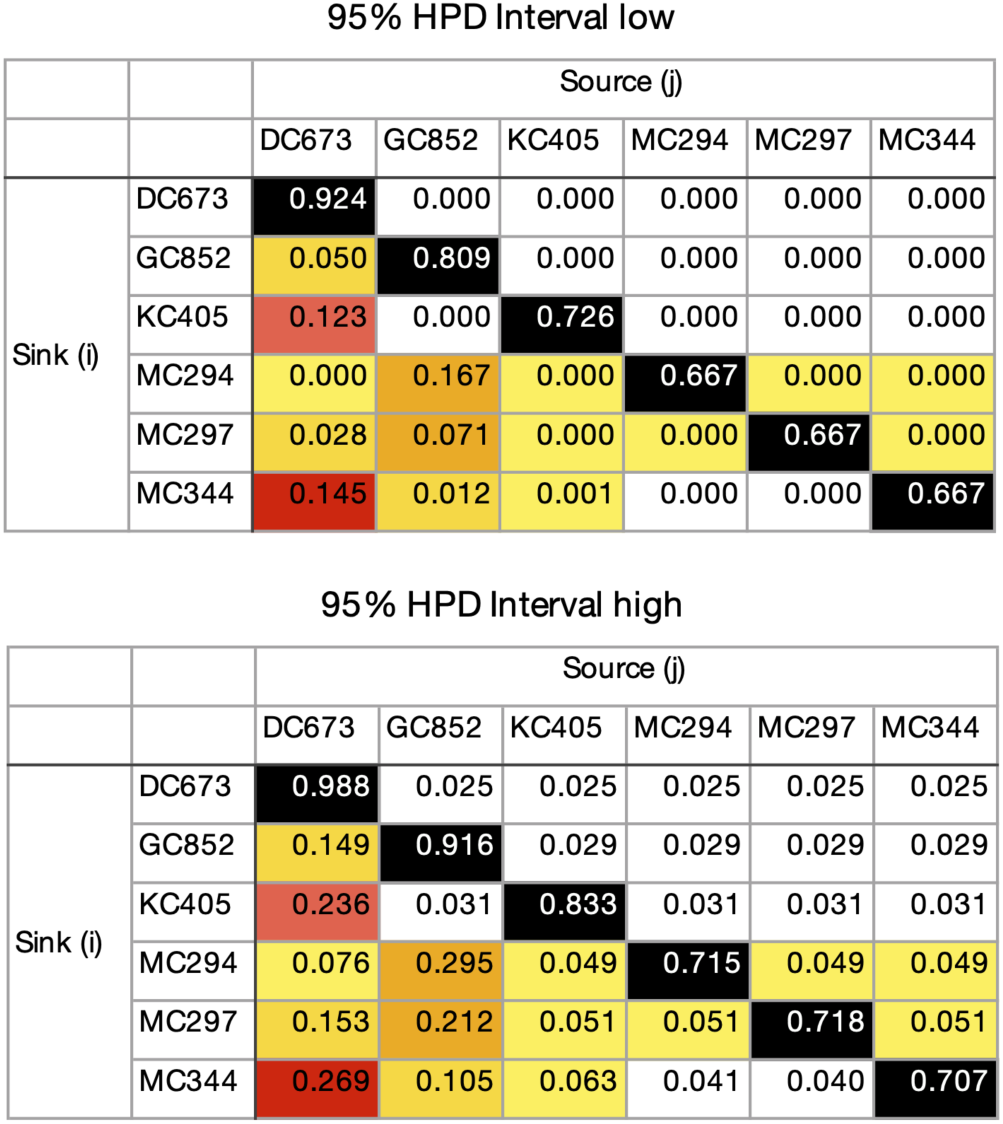
**95%** HPD confidence intervals of migration rates (*m*) estimated in BayesAss

**Figure S3.**
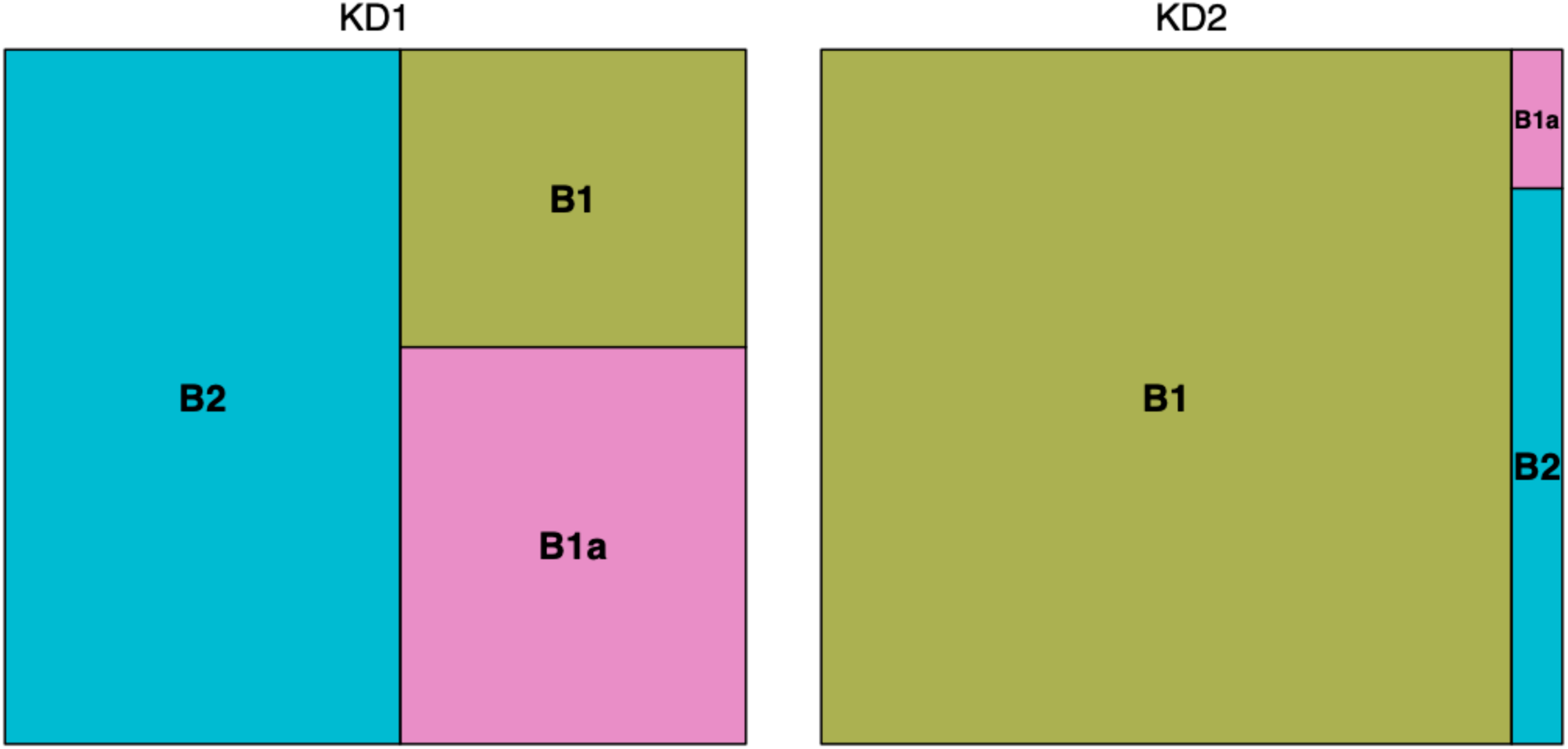
Treemap showing the correspondence between. mitochondrial haplotypes and genomic clusters identified by DAPC analyses (left: K_D1_ and right: K_D2_). The size of each rectangle is proportional to the number of individuals with a given mitochondrial haplotype.

## References

Almany, G. R., Connolly, S. R., Heath, D. D., Hogan, J. D., Jones, G. P., McCook, L. J., et al. (2009). Connectivity, biodiversity conservation and the design of marine reserve networks for coral reefs. Coral Reefs 28, 339–351. doi:10.1007/s00338-009-0484-x.

Amaro, T., Huvenne, V. A. I., Allcock, A. L., Aslam, T., Davies, J. S., Danovaro, R., et al. (2016). The Whittard Canyon – A case study of submarine canyon processes. Prog. Oceanogr. 146, 38–57. doi:10.1016/j.pocean.2016.06.003.

Baco, A. R., Etter, R. J., Ribeiro, P. A., von der Heyden, S., Beerli, P., and Kinlan, B. P. (2016). A synthesis of genetic connectivity in deep-sea fauna and implications for marine reserve design. Mol. Ecol. 25, 3276–3298. doi:10.1111/mec.13689.

Baird, N. A., Etter, P. D., Atwood, T. S., Currey, M. C., Shiver, A. L., Lewis, Z. A., et al. (2008). Rapid SNP discovery and genetic mapping using sequenced RAD markers. PLoS One 3, e3376. doi:10.1371/journal.pone.0003376.

Baums, I. B., Baker, A. C., Davies, S. W., Grottoli, A. G., Kenkel, C. D., Kitchen, S. A., et al. (2019). Considerations for maximizing the adaptive potential of restored coral populations in the western Atlantic. Ecol. Appl. 29, e01978. doi:10.1002/eap.1978.

Behrenfeld, M. J., and Falkowski, P. G. (1997). Photosynthetic rates derived from satellite-based chlorophyll concentration. Limnol. Oceanogr. 42, 1–20. doi:10.4319/lo.1997.42.1.0001.

Benestan, L., Quinn, B. K., Maaroufi, H., Laporte, M., Clark, F. K., Greenwood, S. J., et al. (2016). Seascape genomics provides evidence for thermal adaptation and current-mediated population structure in American lobster (Homarus americanus). Mol. Ecol. 25, 5073–5092. doi:10.1111/mec.13811.

Bernatchez, S., Xuereb, A., Laporte, M., Benestan, L., Steeves, R., Laflamme, M., et al. (2019). Seascape genomics of eastern oyster (Crassostrea virginica) along the Atlantic coast of Canada. Evol. Appl. 12, 587–609. doi:10.1111/eva.12741.

Bertola, L. D., Boehm, J. T., Putman, N. F., Xue, A. T., Robinson, J. D., Harris, S., et al. (2020). Asymmetrical gene flow in five co-distributed syngnathids explained by ocean currents and rafting propensity. Proc. Biol. Sci. 287, 20200657. doi:10.1098/rspb.2020.0657.

Biggs, D. C., Hu, C., and Müller-Karger, F. E. (2008). Remotely sensed sea-surface chlorophyll and POC flux at Deep Gulf of Mexico Benthos sampling stations. Deep Sea Res. Part 2 Top. Stud. Oceanogr. 55, 2555–2562. doi:10.1016/j.dsr2.2008.07.013.

Botsford, L. W., White, J. W., Coffroth, M.-. A., Paris, C. B., Planes, S., Shearer, T. L., et al. (2009). Connectivity and resilience of coral reef metapopulations in marine protected areas: matching empirical efforts to predictive needs. Coral Reefs 28, 327–337. doi:10.1007/s00338-009-0466-z.

Bracco, A., Choi, J., Joshi, K., Luo, H., and McWilliams, J. C. (2016). Submesoscale currents in the northern Gulf of Mexico: Deep phenomena and dispersion over the continental slope. Ocean Model. 101, 43–58. doi:10.1016/j.ocemod.2016.03.002.

Bracco, A., Liu, G., Galaska, M. P., Quattrini, A. M., and Herrera, S. (2019). Integrating physical circulation models and genetic approaches to investigate population connectivity in deep-sea corals. J. Mar. Syst. 198, 103189.

Breusing, C., Biastoch, A., Drews, A., Metaxas, A., Jollivet, D., Vrijenhoek, R. C., et al. (2016). Biophysical and Population Genetic Models Predict the Presence of “Phantom” Stepping Stones Connecting Mid-Atlantic Ridge Vent Ecosystems. Curr. Biol. 26, 2257–2267. doi:10.1016/j.cub.2016.06.062.

Carlon, D. B., and Lippé, C. (2011). Estimation of mating systems in Short and Tall ecomorphs of the coral Favia fragum. Mol. Ecol. 20, 812–828. doi:10.1111/j.1365-294X.2010.04983.x.

Catchen, J., Hohenlohe, P. A., Bassham, S., Amores, A., and Cresko, W. A. (2013). Stacks: an analysis tool set for population genomics. Mol. Ecol. 22, 3124–3140. doi:10.1111/mec.12354.

Cerca, J., Rivera-Colón, A. G., Ferreira, M. S., Ravinet, M., Nowak, M. D., Catchen, J. M., et al. (2021). Incomplete lineage sorting and ancient admixture, and speciation without morphological change in ghost-worm cryptic species. PeerJ 9, e10896. doi:10.7717/peerj.10896.

Cordes, E. E., McGinley, M. P., Podowski, E. L., Becker, E. L., Lessard-Pilon, S., Viada, S. T., et al. (2008). Coral communities of the deep Gulf of Mexico. Deep Sea Res. Part I 55, 777–787. doi:10.1016/j.dsr.2008.03.005.

Danecek, P., Auton, A., Abecasis, G., Albers, C. A., Banks, E., DePristo, M. A., et al. (2011). The variant call format and VCFtools. Bioinformatics 27, 2156–2158. doi:10.1093/bioinformatics/btr330.

Deepwater Horizon Natural Resource Damage Assessment Trustees (2016). Deepwater Horizon oil spill: Final Programmatic Damage Assessment and Restoration Plan and Final Programmatic Environmental Impact Statement. Available at: https://www.gulfspillrestoration.noaa.gov/restoration-planning/gulf-plan [Accessed September 5, 2021].

DeLeo, D. M., Herrera, S., and Lengyel, S. D. (2018). Gene expression profiling reveals deep-sea coral response to the Deepwater Horizon oil spill. Molecular. Available at: https://onlinelibrary.wiley.com/doi/abs/10.1111/mec.14847.

Demopoulos, A. W. J., Bourque, J. R., and Frometa, J. (2014). Biodiversity and community composition of sediment macrofauna associated with deep-sea Lophelia pertusa habitats in the Gulf of Mexico. Deep Sea Res. Part I 93, 91–103. doi:10.1016/j.dsr.2014.07.014.

Dormann, C. F., Elith, J., Bacher, S., Buchmann, C., Carl, G., Carré, G., et al. (2013). Collinearity: a review of methods to deal with it and a simulation study evaluating their performance. Ecography 36, 27–46. doi:10.1111/j.1600-0587.2012.07348.x.

Doughty, C. L., Quattrini, A. M., and Cordes, E. E. (2014). Insights into the population dynamics of the deep-sea coral genus Paramuricea in the Gulf of Mexico. Deep Sea Res. Part 2 Top. Stud. Oceanogr. 99, 71–82. doi:10.1016/j.dsr2.2013.05.023.

Dray, S., Blanchet, G., Borcard, D., Guenard, G., Jombart, T., Larocque, G., et al. (2018). Package “adespatial.” R Package 2018, 3–8. Available at: https://cran.microsoft.com/web/packages/adespatial/adespatial.pdf.

Dray, S., Legendre, P., and Peres-Neto, P. R. (2006). Spatial modelling: a comprehensive framework for principal coordinate analysis of neighbour matrices (PCNM). Ecol. Modell. 196, 483–493. doi:10.1016/j.ecolmodel.2006.02.015.

Dullo W, C., Flögel, S., and Rüggeberg, A. (2008). Cold-water coral growth in relation to the hydrography of the Celtic and Nordic European continental margin. Mar. Ecol. Prog. Ser. 371, 165–176. doi:10.3354/meps07623.

Earl, D. A., and vonHoldt, B. M. (2012). STRUCTURE HARVESTER: a website and program for visualizing STRUCTURE output and implementing the Evanno method. Conserv. Genet. Resour. 4, 359–361. doi:10.1007/s12686-011-9548-7.

Eaton, D. A. R. (2014). PyRAD: assembly of de novo RADseq loci for phylogenetic analyses. Bioinformatics 30, 1844–1849. doi:10.1093/bioinformatics/btu121.

Evanno, G., Regnaut, S., and Goudet, J. (2005). Detecting the number of clusters of individuals using the software STRUCTURE: a simulation study. Mol. Ecol. 14, 2611–2620. doi:10.1111/j.1365-294X.2005.02553.x.

Excoffier, L., Smouse, P. E., and Quattro, J. M. (1992). Analysis of molecular variance inferred from metric distances among DNA haplotypes: application to human mitochondrial DNA restriction data. Genetics 131, 479–491. Available at: https://www.ncbi.nlm.nih.gov/pubmed/1644282.

Fisher, C. R., Hsing, P.-Y., Kaiser, C. L., Yoerger, D. R., Roberts, H. H., Shedd, W. W., et al. (2014). Footprint of Deepwater Horizon blowout impact to deep-water coral communities. Proc. Natl. Acad. Sci. U. S. A. 111, 11744–11749. doi:10.1073/pnas.1403492111.

Foll, M., and Gaggiotti, O. (2008). A genome-scan method to identify selected loci appropriate for both dominant and codominant markers: a Bayesian perspective. Genetics 180, 977–993. doi:10.1534/genetics.108.092221.

Gaines, S. D., White, C., Carr, M. H., and Palumbi, S. R. (2010). Designing marine reserve networks for both conservation and fisheries management. Proc. Natl. Acad. Sci. U. S. A. 107, 18286–18293. doi:10.1073/pnas.0906473107.

Gaither, M. R., Gkafas, G. A., de Jong, M., Sarigol, F., Neat, F., Regnier, T., et al. (2018). Genomics of habitat choice and adaptive evolution in a deep-sea fish. Nat Ecol Evol 2, 680–687. doi:10.1038/s41559-018-0482-x.

Galaska, M. P., Sands, C. J., Santos, S. R., Mahon, A. R., and Halanych, K. M. (2017). Crossing the Divide: Admixture Across the Antarctic Polar Front Revealed by the Brittle Star Astrotoma agassizii. Biol. Bull. 232, 198–211. doi:10.1086/693460.

Galindo, H. M., Olson, D. B., and Palumbi, S. R. (2006). Seascape Genetics: A Coupled Oceanographic-Genetic Model Predicts Population Structure of Caribbean Corals. Curr. Biol. 16, 1622–1626. doi:10.1016/j.cub.2006.06.052.

Gan, Z., Yuan, J., Liu, X., Dong, D., Li, F., and Li, X. (2020). Comparative transcriptomic analysis of deep-and shallow-water barnacle species (Cirripedia, Poecilasmatidae) provides insights into deep-sea adaptation of sessile crustaceans. BMC Genomics 21, 240. doi:10.1186/s12864-020-6642-9.

Georgian, S. E., Kramer, K., Saunders, M., Shedd, W., Roberts, H., Lewis, C., et al. (2020). Habitat suitability modelling to predict the spatial distribution of cold-water coral communities affected by the Deepwater Horizon oil spill. J. Biogeogr. 47, 1455–1466. doi:10.1111/jbi.13844.

Girard, F., Cruz, R., Glickman, O., Harpster, T., Fisher, C. R., and Thomsen, L. (2019). In situ growth of deep-sea octocorals after the Deepwater Horizon oil spill. Elementa: Science of the Anthropocene 7. doi:10.1525/journal.elementa.349.

González, A. M., Prada, C. A., Ávila, V., and Medina, M. (2018). “Ecological Speciation in Corals,” in Population Genomics (Cham: Springer International Publishing), 303–324. doi:10.1007/13836_2018_35.

Gosselin, T. assigner. Github Available at: https://github.com/thierrygosselin/assigner [Accessed May 13, 2021].

Goudet, J. (2005). hierfstat, a package for r to compute and test hierarchical F-statistics. Mol. Ecol. Notes 5, 184–186. doi:10.1111/j.1471-8286.2004.00828.x.

Goyert, H. F., Bassett, R., Christensen, J., Coleman, H, Coyne, M., et al. Characterizing spatial distributions of deep-sea corals and chemosynthetic communities in the US Gulf of Mexico through data synthesis and predictive modeling. US Department of the Interior, Bureau of Ocean Energy Management Available at: https://espis.boem.gov/final%20reports/BOEM_2021-027.pdf.

Hellberg, M. E. (2009). Gene Flow and Isolation among Populations of Marine Animals. Annu. Rev. Ecol. Evol. Syst. 40, 291–310. doi:10.1146/annurev.ecolsys.110308.120223.

Henry, L.-A., and Roberts, J. M. (2007). Biodiversity and ecological composition of macrobenthos on cold-water coral mounds and adjacent off-mound habitat in the bathyal Porcupine Seabight, NE Atlantic. Deep Sea Res. Part I 54, 654–672. doi:10.1016/j.dsr.2007.01.005.

Herrera, S., Reyes-Herrera, P. H., and Shank, T. M. (2015). Predicting RAD-seq Marker Numbers across the Eukaryotic Tree of Life. Genome Biol. Evol. 7, 3207–3225. doi:10.1093/gbe/evv210.

Herrera, S., and Shank, T. M. (2016). RAD sequencing enables unprecedented phylogenetic resolution and objective species delimitation in recalcitrant divergent taxa. Mol. Phylogenet. Evol. 100, 70–79.

Hilario, A., Metaxas, A., Gaudron, S. M., Howell, K. L., Mercier, A., Mestre, N. C., et al. (2015). Estimating dispersal distance in the deep sea: challenges and applications to marine reserves. Front. Mar. Sci. 2. doi:10.3389/fmars.2015.00006.

Hoffman, J. I., Clarke, A., Clark, M. S., Fretwell, P., and Peck, L. S. (2012). Unexpected fine-scale population structure in a broadcast-spawning Antarctic marine mollusc. PLoS One 7, e32415. doi:10.1371/journal.pone.0032415.

Jombart, T. (2008). adegenet: a R package for the multivariate analysis of genetic markers. Bioinformatics 24, 1403–1405. doi:10.1093/bioinformatics/btn129.

Jones, G. P., Srinivasan, M., and Almany, G. R. (2007). Population Connectivity and Conservation of Marine Biodiversity. Oceanography 20, 100–111. Available at: http://www.jstor.org/stable/24860100.

Kamvar, Z. N., Tabima, J. F., and Grünwald, N. J. (2014). Poppr: an R package for genetic analysis of populations with clonal, partially clonal, and/or sexual reproduction. PeerJ 2, e281. doi:10.7717/peerj.281.

Kelley, D. E. (2018). “The oce Package,” in Oceanographic Analysis with R, ed. D. E. Kelley (New York, NY: Springer New York), 91–101. doi:10.1007/978-1-4939-8844-0_3.

Kenchington, E., Yashayaev, I., Tendal, O. S., and Jørgensbye, H. (2017). Water mass characteristics and associated fauna of a recently discovered Lophelia pertusa (Scleractinia: Anthozoa) reef in Greenlandic waters. Polar Biol. 40, 321–337. doi:10.1007/s00300-016-1957-3.

Kinlan, B. P., Gaines, S. D., and Lester, S. E. (2005). Propagule dispersal and the scales of marine community process. Divers. Distrib. 11, 139–148. doi:10.1111/j.1366-9516.2005.00158.x.

Lan, Y., Sun, J., Tian, R., Bartlett, D. H., Li, R., Wong, Y. H., et al. (2017). Molecular adaptation in the world’s deepest-living animal: Insights from transcriptome sequencing of the hadal amphipod Hirondellea gigas. Mol. Ecol. 26, 3732–3743. doi:10.1111/mec.14149.

Lan, Y., Sun, J., Xu, T., Chen, C., Tian, R., Qiu, J.-W., et al. (2018). De novo transcriptome assembly and positive selection analysis of an individual deep-sea fish. BMC Genomics 19, 394. doi:10.1186/s12864-018-4720-z.

Legendre, P., and Legendre, L. (2012). Numerical Ecology. Elsevier Available at: https://www.elsevier.com/books/numerical-ecology/legendre/978-0-444-53868-0 [Accessed September 9, 2021].

Lemaire, B., Karchner, S. I., Goldstone, J. V., Lamb, D. C., Drazen, J. C., Rees, J. F., et al. (2018). Molecular adaptation to high pressure in cytochrome P450 1A and aryl hydrocarbon receptor systems of the deep-sea fish Coryphaenoides armatus. Biochim. Biophys. Acta: Proteins Proteomics 1866, 155–165. doi:10.1016/j.bbapap.2017.06.026.

Lett, C., Verley, P., Mullon, C., Parada, C., Brochier, T., Penven, P., et al. (2008). A Lagrangian tool for modelling ichthyoplankton dynamics. Environmental Modelling & Software 23, 1210–1214. doi:10.1016/j.envsoft.2008.02.005.

Lipcius, R. N., Eggleston, D. B., Schreiber, S. J., Seitz, R. D., Shen, J., Sisson, M., et al. (2008). Importance of Metapopulation Connectivity to Restocking and Restoration of Marine Species. Rev. Fish. Sci. 16, 101–110. doi:10.1080/10641260701812574.

Liu, G., Bracco, A., Quattrini, A. M., and Herrera, S. Kilometer-scale larval dispersal processes and the connectivity of Paramuricea biscaya in the northern Gulf of Mexico.

Maiti, K., Bosu, S., D’Sa, E. J., Adhikari, P. L., Sutor, M., and Longnecker, K. (2016). Export fluxes in northern Gulf of Mexico - Comparative evaluation of direct, indirect and satellite-based estimates. Mar. Chem. 184, 60–77. doi:10.1016/j.marchem.2016.06.001.

Martin, J. H., Knauer, G. A., Karl, D. M., and Broenkow, W. W. (1987). VERTEX: carbon cycling in the northeast Pacific. Deep Sea Res. A 34, 267–285. doi:10.1016/0198-0149(87)90086-0.

McArdle, B. H., and Anderson, M. J. (2001). Fitting multivariate models to community data: A comment on distance-based redundancy analysis. Ecology 82, 290–297. doi:10.1890/0012-9658(2001)082[0290:fmmtcd]2.0.co;2.

McClain, C. R., and Mincks Hardy, S. (2010). The biogeography of the deep sea: range and dispersal on the seafloor. Proc. Roy. Soc. B-Biol Sci.

Meirmans, P. G. (2020). genodive version 3.0: Easy-to-use software for the analysis of genetic data of diploids and polyploids. Mol. Ecol. Resour. 20, 1126–1131. doi:10.1111/1755-0998.13145.

Miller, K. J., Rowden, A. A., Williams, A., and Häussermann, V. (2011). Out of their depth? Isolated deep populations of the cosmopolitan coral Desmophyllum dianthus may be highly vulnerable to environmental change. PLoS One 6, e19004. doi:10.1371/journal.pone.0019004.

Mouw, C. B., Barnett, A., McKinley, G. A., Gloege, L., and Pilcher, D. (2016). Global Ocean Particulate Organic Carbon flux merged with satellite parameters, supplement to: Mouw, Colleen B; Barnett, Audrey; McKinley, Galen A; Gloege, Lucas; Pilcher, Darren (2016): Global ocean particulate organic carbon flux merged with satellite parameters. Earth System Science Data, 8(2), 531–541. doi:10.1594/PANGAEA.855600.

Mullineaux, L. S., Metaxas, A., Beaulieu, S. E., Bright, M., Gollner, S., Grupe, B. M., et al. (2018). Exploring the ecology of deep-sea hydrothermal vents in a metacommunity framework. Frontiers in Marine Science 5, 49.

Nei, M. (1978). Estimation of average heterozygosity and genetic distance from a small number of individuals. Genetics 89, 583–590. Available at: https://www.ncbi.nlm.nih.gov/pubmed/17248844.

Oksanen, J., Kindt, R., Legendre, P., O’Hara, B., Stevens, M. H. H., Oksanen, M. J., et al. (2007). The vegan package. Community ecology package 10, 719. Available at: https://www.researchgate.net/profile/Gavin-Simpson-2/publication/228339454_The_vegan_Package/links/0912f50be86bc29a7f000000/The-vegan-Package.pdf.

Pace, M. L., Knauer, G. A., Karl, D. M., and Martin, J. H. (1987). Primary production, new production and vertical flux in the eastern Pacific Ocean. Nature 325, 803–804. doi:10.1038/325803a0.

Palumbi, S. R. (2003). Population genetics, demographic connectivity, and the design of marine reserves. Ecol. Appl. 13, 146–158. doi:10.1890/1051-0761(2003)013[0146:pgdcat]2.0.co;2.

Pante, E., Abdelkrim, J., Viricel, A., Gey, D., France, S. C., Boisselier, M. C., et al. (2015). Use of RAD sequencing for delimiting species. Heredity 114, 450–459. doi:10.1038/hdy.2014.105.

Pante, E., and Simon-Bouhet, B. (2013). marmap: A package for importing, plotting and analyzing bathymetric and topographic data in R. PLoS One 8, e73051. doi:10.1371/journal.pone.0073051.

Paradis, E. (2010). pegas: an R package for population genetics with an integrated–modular approach. Bioinformatics 26, 419–420. doi:10.1093/bioinformatics/btp696.

Parra-Salazar, A., Gomez, J., Lozano-Arce, D., Reyes-Herrera, P. H., and Duitama, J. (2021). Robust and efficient software for reference-free genomic diversity analysis of genotyping-by-sequencing data on diploid and polyploid species. Mol. Ecol. Resour. doi:10.1111/1755-0998.13477.

Pritchard, J. K., Stephens, M., and Donnelly, P. (2000). Inference of population structure using multilocus genotype data. Genetics 155, 945–959. Available at: https://www.ncbi.nlm.nih.gov/pubmed/10835412.

Prouty, N. G., Fisher, C. R., Demopoulos, A. W. J., and Druffel, E. R. M. (2016). Growth rates and ages of deep-sea corals impacted by the Deepwater Horizon oil spill. Deep Sea Res. Part 2 Top. Stud. Oceanogr. 129, 196–212. doi:10.1016/j.dsr2.2014.10.021.

Prouty, N. G., Roark, E. B., Buster, N. A., and Ross, S. W. (2011). Growth rate and age distribution of deep-sea black corals in the Gulf of Mexico. Mar. Ecol. Prog. Ser. 423, 101–115. doi:10.3354/meps08953.

Puckett, B. J., and Eggleston, D. B. (2016). Metapopulation dynamics guide marine reserve design: importance of connectivity, demographics, and stock enhancement. Ecosphere 7, e01322. doi:10.1002/ecs2.1322.

Quattrini, A. M., Baums, I. B., Shank, T. M., Morrison, C. L., and Cordes, E. E. (2015). Testing the depth-differentiation hypothesis in a deepwater octocoral. Proc. Biol. Sci. 282, 20150008. doi:10.1098/rspb.2015.0008.

Quattrini, A. M., Etnoyer, P. J., Doughty, C., English, L., Falco, R., Remon, N., et al. (2014). A phylogenetic approach to octocoral community structure in the deep Gulf of Mexico. Deep Sea Res. Part 2 Top. Stud. Oceanogr. 99, 92–102. doi:10.1016/j.dsr2.2013.05.027.

Quattrini, A. M., Gómez, C. E., and Cordes, E. E. (2017). Environmental filtering and neutral processes shape octocoral community assembly in the deep sea. Oecologia 183, 221–236. doi:10.1007/s00442-016-3765-4.

Quattrini, A. M., Wu, T., Soong, K., Jeng, M.-S., Benayahu, Y., and McFadden, C. S. (2019). A next generation approach to species delimitation reveals the role of hybridization in a cryptic species complex of corals. BMC Evol. Biol. 19, 116. doi:10.1186/s12862-019-1427-y.

Radice, V. Z., Quattrini, A. M., Wareham, V. E., Edinger, E. N., and Cordes, E. E. (2016). Vertical water mass structure in the North Atlantic influences the bathymetric distribution of species in the deep-sea coral genus Paramuricea. Deep Sea Res. Part I 116, 253–263. doi:10.1016/j.dsr.2016.08.014.

Rambaut, A., Drummond, A. J., Xie, D., Baele, G., and Suchard, M. A. (2018). Posterior Summarization in Bayesian Phylogenetics Using Tracer 1.7. Syst. Biol. 67, 901–904. doi:10.1093/sysbio/syy032.

Reitzel, A. M., Herrera, S., Layden, M. J., Martindale, M. Q., and Shank, T. M. (2013). Going where traditional markers have not gone before: utility of and promise for RAD sequencing in marine invertebrate phylogeography and population genomics. Mol. Ecol. 22, 2953–2970. doi:10.1111/mec.12228.

Roark, E. B., Guilderson, T. P., Dunbar, R. B., Fallon, S. J., and Mucciarone, D. A. (2009). Extreme longevity in proteinaceous deep-sea corals. Proc. Natl. Acad. Sci. U. S. A. 106, 5204–5208. doi:10.1073/pnas.0810875106.

Roberts, E. M., Bowers, D. G., Meyer, H. K., Samuelsen, A., Rapp, H. T., and Cárdenas, P. (2021). Water masses constrain the distribution of deep-sea sponges in the North Atlantic Ocean and Nordic Seas. Mar. Ecol. Prog. Ser. 659, 75–96. doi:10.3354/meps13570.

Ross, S. W., and Quattrini, A. M. (2007). The fish fauna associated with deep coral banks off the southeastern United States. Deep Sea Res. Part I 54, 975–1007. doi:10.1016/j.dsr.2007.03.010.

Rowden, A. A., Dower, J. F., Schlacher, T. A., Consalvey, M., and Clark, M. R. (2010). Paradigms in seamount ecology: fact, fiction and future. Mar. Ecol. 31, 226–241. doi:10.1111/j.1439-0485.2010.00400.x.

Sandoval-Castillo, J., Robinson, N. A., Hart, A. M., Strain, L. W. S., and Beheregaray, L. B. (2018). Seascape genomics reveals adaptive divergence in a connected and commercially important mollusc, the greenlip abalone (Haliotis laevigata), along a longitudinal environmental gradient. Mol. Ecol. 27, 1603–1620. doi:10.1111/mec.14526.

Selkoe, K. A., D’Aloia, C. C., Crandall, E. D., Iacchei, M., Liggins, L., Puritz, J. B., et al. (2016). A decade of seascape genetics: contributions to basic and applied marine connectivity. Mar. Ecol. Prog. Ser. 554, 1–19. doi:10.3354/meps11792.

Serrano, X. M., Baums, I. B., Smith, T. B., Jones, R. J., Shearer, T. L., and Baker, A. C. (2016). Long distance dispersal and vertical gene flow in the Caribbean brooding coral Porites astreoides. Sci. Rep. 6, 21619. doi:10.1038/srep21619.

Sherwood, O. A., and Edinger, E. N. (2009). Ages and growth rates of some deep-sea gorgonian and antipatharian corals of Newfoundland and Labrador. Can. J. Fish. Aquat. Sci. 66, 142–152. doi:10.1139/f08-195.

Smith, K. L., Jr, Ruhl, H. A., Huffard, C. L., Messié, M., and Kahru, M. (2018). Episodic organic carbon fluxes from surface ocean to abyssal depths during long-term monitoring in NE Pacific. Proc. Natl. Acad. Sci. U. S. A. 115, 12235–12240. doi:10.1073/pnas.1814559115.

Somero, G. N. (1992). Adaptations to high hydrostatic pressure. Annu. Rev. Physiol. 54, 557–577. doi:10.1146/annurev.ph.54.030192.003013.

Taylor, M. L., and Roterman, C. N. (2017). Invertebrate population genetics across Earth’s largest habitat: The deep-sea floor. Mol. Ecol. 26, 4872–4896. doi:10.1111/mec.14237.

Team, R. C., and Others (2013). R: A language and environment for statistical computing. Available at: http://r.meteo.uni.wroc.pl/web/packages/dplR/vignettes/intro-dplR.pdf.

Tello, D., Gil, J., Loaiza, C. D., Riascos, J. J., Cardozo, N., and Duitama, J. (2019). NGSEP3: accurate variant calling across species and sequencing protocols. Bioinformatics 35, 4716–4723. doi:10.1093/bioinformatics/btz275.

Tonkin-Hill, G., and Lee, S. (2016). starmie: Population Structure Model Inference and Visualisation. Available at: https://cran.r-proje.

Vohsen, S. A., Anderson, K. E., Gade, A. M., Gruber-Vodicka, H. R., Dannenberg, R. P., Osman, E. O., et al. (2020). Deep-sea corals provide new insight into the ecology, evolution, and the role of plastids in widespread apicomplexan symbionts of anthozoans. Microbiome 8, 34. doi:10.1186/s40168-020-00798-w.

Vurture, G. W., Sedlazeck, F. J., Nattestad, M., Underwood, C. J., Fang, H., Gurtowski, J., et al. (2017). GenomeScope: fast reference-free genome profiling from short reads. Bioinformatics 33, 2202–2204. doi:10.1093/bioinformatics/btx153.

Weber, A. A.-T., Hugall, A. F., and O’Hara, T. D. (2020). Convergent Evolution and Structural Adaptation to the Deep Ocean in the Protein-Folding Chaperonin CCTα. Genome Biol. Evol. 12, 1929–1942. doi:10.1093/gbe/evaa167.

Weir, B. S., and Cockerham, C. C. (1984). ESTIMATING F-STATISTICS FOR THE ANALYSIS OF POPULATION STRUCTURE. Evolution 38, 1358–1370. doi:10.1111/j.1558-5646.1984.tb05657.x.

White, H. K., Hsing, P.-Y., Cho, W., Shank, T. M., Cordes, E. E., Quattrini, A. M., et al. (2012). Impact of the Deepwater Horizon oil spill on a deep-water coral community in the Gulf of Mexico. Proc. Natl. Acad. Sci. U. S. A. 109, 20303–20308. doi:10.1073/pnas.1118029109.

Wilson, A. M., Raine, R., Mohn, C., and White, M. (2015). Nepheloid layer distribution in the Whittard Canyon, NE Atlantic Margin. Mar. Geol. 367, 130–142. doi:10.1016/j.margeo.2015.06.002.

Wilson, G. A., and Rannala, B. (2003). Bayesian inference of recent migration rates using multilocus genotypes. Genetics 163, 1177–1191. Available at: https://www.ncbi.nlm.nih.gov/pubmed/12663554.

Woolley, S. N. C., Tittensor, D. P., Dunstan, P. K., Guillera-Arroita, G., Lahoz-Monfort, J. J., Wintle, B. A., et al. (2016). Deep-sea diversity patterns are shaped by energy availability. Nature 533, 393–396. doi:10.1038/nature17937.

Xuereb, A., Benestan, L., Normandeau, É., Daigle, R. M., Curtis, J. M. R., Bernatchez, L., et al. (2018). Asymmetric oceanographic processes mediate connectivity and population genetic structure, as revealed by RADseq, in a highly dispersive marine invertebrate (Parastichopus californicus). Mol. Ecol. 27, 2347–2364. doi:10.1111/mec.14589.

Zeng, C., Rowden, A. A., Clark, M. R., and Gardner, J. P. A. (2020). Species-specific genetic variation in response to deep-sea environmental variation amongst Vulnerable Marine Ecosystem indicator taxa. Sci. Rep. 10, 2844. doi:10.1038/s41598-020-59210-0.

